# New structures of Class II Fructose-1,6-Bisphosphatase from *Francisella tularensis* suggest a novel catalytic mechanism for the entire Class

**DOI:** 10.1101/2022.09.05.506711

**Authors:** Anna I. Selezneva, Luke N. M. Harding, Hiten J. Gutka, Farahnaz Movahedzadeh, Celerino Abad-Zapatero

## Abstract

Class II Fructose-1,6-bisphosphatases (FBPaseII) (EC: 3.1.3.11) are highly conserved essential enzymes in the gluconeogenic pathway of microorganisms. Previous crystallographic studies of FBPasesII provided insights into various inactivated states of enzymes from different species. Presented here is the first crystal structure of FBPaseII in an active state, solved for the enzyme from *Francisella tularensis* (*Ft*FBPaseII), containing native metal cofactor Mn^2+^ and complexed with catalytic product fructose-6-phosphate (F6P). Another crystal structure of the same enzyme complex is presented in inactivated state due to the structural changes introduced by crystal packing. Analysis of the interatomic distances among the substrate, product and divalent metal cations in the enzyme catalytic centers led to a revision of the catalytic mechanism suggested previously for class II FBPases. Instead of a metal cofactor for the stabilization of the transition state of the leaving phosphate group, we propose that the positive dipole of the neighboring α-helix backbone (**G88-T89-T90-I91-T92-S93-K94**) is responsible for retaining the cleaved phosphate. The revised catalytic mechanism involves a nucleophilic attack by a reactive water coordinated by both the T89 hydroxyl and a second water molecule coordinated directly by Mn^2+^. Additionally, a crystal structure of *Mycobacterium tuberculosis* FBPaseII (*Mt*FBPaseII), containing the substrate fructose-1,6-bisphosphate (F1,6BP) bound, is presented in support of a novel catalytic mechanism for this class of enzymes.

## Introduction

Gluconeogenesis is a critical metabolic pathway for the survival of organisms when dietary intake of glucose is insufficient or absent. Class II fructose-1,6-bisphosphatase (FBPaseII, EC: 3.1.3.11) has been shown to be one of the key gluconeogenesis enzymes that is essential for the virulence of *Francisella tularensis* [1], a highly pathogenic Gram-negative coccobacillus species that causes tularemia in humans around the world. Due to the potential lethality and rarity of tularemia in mainstream medicine, *F. tularensis* has been recognized as a category A bioterrorism agent [2]. The pathogen replicates intracellularly to high densities mainly in human macrophages [3], utilizing host-derived fatty acids, pyruvate, glycerol and amino acids as a carbon, nitrogen, and energy source for the *de novo* synthesis of molecules that are essential for the pathogen’s survival and growth. *F. tularensis* FBPaseII (*Ft*FBPaseII) is encoded by the *glpX* gene, and catalyzes the hydrolysis of F1,6BP to F6P and an inorganic phosphate. Its product F6P is a precursor of the pentose phosphate pathway and necessary in the *de novo* synthesis of pentose phosphates, lipopolysaccharides, and peptidoglycan — all molecules that are crucial for the pathogen’s intracellular survival.

FBPases are a diverse group of enzymes, consisting of five structurally different classes (I-V) sharing similarities in the overall domain organization. All FBPases possess two adjacent α/β domains, one of which is two-layered (α-β; CATH code 3.30.540), and the other is three-layered (α-β-α; CATH code 3.40.190). The two domains combine to form a five-layered sandwich-like structure in which the β-sheets are positioned nearly orthogonal to each other. Class-specific differences are determined by the number of strands in each sheet, the boundaries of the strands and the topology connecting them. Class II FBPases are highly conserved enzymes with over 100 species sharing 50% or higher sequence identity, whereas only about 10% of sequence identity is shared with FBPases of other classes [4]. Prokaryotic FBPases include members of all five classes, while eukaryotic FBPases are mostly limited to class I. This and the fact that no other FBPases have been discovered in *F. tularensis* to this day, makes *Ft*FBPaseII an attractive target for the structure-guided drug design of potentially bactericidal FBPaseII inhibitors innocuous to mammalian enzymes (selective to class II enzymes).

Despite the growing number of structures available for class II FBPases, the details of the catalytic mechanism and the distinctions determining metal dependence of the members of the class remain under-investigated. Previous attempts to investigate a catalytic mechanism of FBPaseII involved crystal structures of *Escherichia coli* FBPaseII (*Ec*FBPaseII) [4]; PDB entries 3big, 3bih and 3d1r. The proposed mechanism was based on the *Ec*FBPaseII D61A mutant crystal structure (PDB entry 3d1r). The structure revealed the presence of two Ca^2+^ ions, one Mg^2+^ ion (coordinated in between the two phosphate groups of the substrate on the opposite site) and a molecule of substrate F1,6BP in the active center of the enzyme. However, biochemical characterization of *Ec*FBPaseII by the same authors demonstrated that not only is mutant D61A essentially inactive, but Mn^2+^ — which is absent from the structure — is solely required for *Ec*FBPaseII activity, not Ca^2+^ nor Mg^2+^. The inactivity observed was consistent with the previous observations claiming that replacement of Mn^2+^ with CaCl_2_ or MgCl_2_ results in almost complete loss of activity [4, 5]. Nevertheless, the authors hypothesized a catalytic mechanism involving two Ca^2+^ ions in the active center: one to stabilize the negative charge on the leaving phosphate group and one to coordinate the nucleophilic water. The authors acknowledged that the Mg^2+^ ion seems not to be involved in catalysis as it is liganded to the inactive phosphate-6 in the substrate molecule. Hence, the available structure of *Ec*FBPaseII D61A with Mg^2+^ and Ca^2+^ in the active center is not a representation of the active state of the enzyme and does not provide appropriate details towards the mechanism of hydrolysis.

Currently, only one published FBPaseII structure contains catalytic product in a combination with appropriate metal cofactor (PDB entry 6ayu). This is a crystal structure of the partially active T84S mutant of *M. tuberculosis* FBPaseII (*Mt*FBPaseII) complexed with Mg^2+^ and F6P. However, the presence of glycerol in the active center makes it difficult to infer a consistent catalytic mechanism. Our own previous attempts to solve the *Ft*FBPaseII structure resulted in the crystal structure of the Mg^2+^-inactivated enzyme [6]; PDB entry 7js3. This conclusion was based in part on the *Ft*FBPaseII activity studies in the presence of different divalent cations [7]. Similar to *Ec*FBPaseII, it was shown that *Ft*FBPaseII depends solely on Mn^2+^ for activity, with Mg^2+^ and Ca^2+^ not supporting the catalysis. However, while Mn^2+^ was examined at a broad concentration range, Mg^2+^ and other ions were studied at a single concentration of 100 mM. It was demonstrated [8] that the amplitude of FBPasesII activity may vary significantly depending on the concentration of the metal cofactor in question, ranging from relatively high enzyme activity to its complete loss. Moreover, in some cases either Mg^2+^ or Mn^2+^ can activate a class II enzyme, while in others only Mn^2+^ is required for activity. Presented in this work is the investigation of *Ft*FBPaseII activity within a broad range of Mn^2+^ and Mg^2+^ concentrations. There is no evidence that class II FBPases require other metals for catalysis besides Mg and Mn. In contrast, FBPases of other classes can be activated by Zn^2+^, for example [9, 10]. Presented in this work is the crystal structure of the wild type *Ft*FBPaseII in its active state containing both F6P and a catalytic metal cofactor Mn^2+^. The structure allows for identification of the residues responsible for coordination of both ligands and for modeling of the missing phosphate group into the active site. Additionally, for the purposes of discussion, we present the inactive state crystal structures of the wild type *Ft*FBPaseII containing F6P and Mn^2+^, and a serendipitous structure of the wild type *Mt*FBPaseII liganded with substrate F1,6BP.

This work focuses on improving our understanding of structure and function of the entire class II FBPases, combining analysis of existing and novel structures together with experimental investigation into the properties of *Mt* and *Ft*FBPasesII and for other important pathogens such as *Staphylococcus aureus*. The results provide support for a novel catalytic mechanism for this class of enzymes, which would involve the positive charge at the N-terminal dipole of a conserved helix, as a critical element to stabilize the transition state of the cleavable phosphate.

## Materials and Methods

### Gene cloning, production, purification of recombinant *Ft*FBPaseII and storage

A previously engineered construct of the *F. tularensis glpX* gene containing an N-terminal His6 tag in pET-15b vector was used for overexpression of the *Ft*FBPaseII enzyme in *E. coli* [7]. Overexpression of the wild-type *Ft*FBPasesII and the purification of the proteins from *E. coli* was performed as described previously [6]. The protein was purified, concentrated to at least 10 mg/ml and stored as single-use aliquots at 193 K until further use in 20 mM Tris–HCl pH 8.0, 50 mM KCl, 10% glycerol. All purification procedures were performed at 277 K.

### Enzymatic assays

Phosphatase activity was quantified spectrophotometrically in Absorbance Units (AU) at 630 nm using colorimetric Malachite Green assay. The reaction mixture (80 µl) contained 40 mM Tris– HCl pH 8.0, 100 mM KCl, 12.5 µg/ml of *Ft*FBPaseII and 450 μM F1,6BP. MgCl_2_ and MnCl_2_ varied as necessary. After 10 min of incubation at 293 K the reaction was quenched by the addition of 20 μl of Malachite Green reagent [11]. Then the color was allowed to develop for an additional 3 min at 293 K. Negative control reactions contained either no metal cofactor or enzyme. Positive control contained alternative FBPasesII. All reagents used for protein purification and enzymatic assays were from Sigma.

### Protein crystallization

#### Crystal form A

Complex of *Ft*FBPaseII with Mn^2+^ and F6P was prepared prior to crystallization. The purified protein sample in 20 mM Tris–HCl pH 8.0, 50 mM KCl, and 10% glycerol was sequentially mixed with solution of MnCl_2_ in 100 mM KCl, and then with solution of F6P in 100 mM KCl. These conditions were chosen to match the ones at which the *Ft*FBPaseII was catalytically active (**Fig. 2B**). The resulting protein concentration in the sample was 11.45 mg/ml, both F6P and MnCl_2_ were at 1 mM each. Precipitant was added immediately upon the complex preparation. Crystals were grown at 291 K using the hanging drop vapor diffusion method [12], combining the enzyme-containing sample with precipitant in a 2:1-sample:precipitant ratio. The crystal used for diffraction studies was obtained with precipitant containing 8% tacsimate pH 6.0 and 20% polyethylene glycol 3,350 from Hampton Research. The crystal was cryoprotected by a solution of 33% glycerol in the crystallization buffer and flash-frozen in liquid nitrogen.

#### Crystal form B

Prior to crystallization, the sample of purified *Ft*FBPaseII in 20 mM Tris–HCl pH 8.0, 50 mM KCl, and 10% glycerol was mixed with 200 mM MnCl_2_ solution in 100 mM KCl and then with 4 mM F1,6BP in 100 mM KCl. The conditions for crystallization were chosen to be near those at which the *Ft*FBPaseII was catalytically active (**Fig. 2B**). The resulting protein concentration in the sample was 10 mg/ml, MnCl_2_ was at 50 mM and F1,6BP was at 1 mM. Precipitant was added immediately upon the addition of substrate. Crystallization was achieved as described above for Form A and a diffractable crystal was obtained in 0.2 M sodium malonate pH 6.0 and 20% polyethylene glycol 3350 from Hampton Research. The crystal was flash-frozen in liquid nitrogen without cryopreservation.

#### Crystal form C

Crystals of *Mt*FBPaseII complexed with the F1,6BP substrate were obtained by mixing 100 μl of protein solution with 10 μl of 10 mM substrate solution. The protein solution contained 1 mM MgCl_2_. These conditions were near the ones at which *Mt*FBPaseII was catalytically inactive (**Fig. 2A**). The sample was incubated for 30 min at 303 K and then hanging droplets were formed by addition of precipitant at 1:1 ratio. Well diffracting crystals were obtained in 2.9 M sodium malonate pH 4.0 (Malonate grid from Hampton Research). The crystals were typically medium size (20 x 30 x 50 μm) and had a well-defined morphology of hexagonal bipyramids.

### Data collection, structure determination and refinement

X-ray diffraction data for both crystal forms A and B were collected at 100 K on the LS-CAT 21-ID-D beamline, using a Geiger 9M detector at the Advanced Photon Source (APS), Argonne National Laboratory, Illinois, USA. All the structures were solved by molecular replacement (MR) using the software MOLREP [13] as implemented in the software suite CCP4 [14]. QtMG [15] was used to produce publication-quality figures.

#### Crystal form A

A wedge of 180° of data were collected in fine slicing mode (0.20° per image, 900) at a crystal to detector distance of 200 mm with an exposure of 0.07 secs per image. The corresponding structure (Form A) was solved by molecular replacement using the tetramer of the Mg^2+^-containing structure (PDB entry 7js3) as a search model. The data were processed and reduced with HKL2000 [16]. The refinement calculations were done using predominantly PHENIX [17] and alternating with the REFMAC5 [18] implementation in CCP4 [14] to facilitate the interpretation of the very external loops that were often smoothed by the bulk solvent correction in PHENIX. Glycerol molecules were added when putative water molecules did not satisfy the positive (3s) F_o_-F_c_ electron density peaks. Structure rebuilding and revisions were done using COOT as implemented in the PHENIX suite [17].

#### Crystal form B

A wedge of 180° of data were collected in fine slicing mode (0.25° per image, 720) at a crystal to detector distance of 250 mm with an exposure of 0.08 secs per image. The data were processed and reduced with HKL2000 [16]. The structure (Form B) was solved by molecular replacement using a partially refined model of Form A, confirming the presence of two *Ft*FBPaseII tetramers in very similar orientations separated by about 70 Å along the crystal c axis. The refinement protocols were analogous to the ones used for Form A.

#### Crystal form C

These crystals were obtained at the earlier stages of the *Mt*FBPaseII project and since they diffracted only to about 3.7 Å resolution they were not fully analyzed at the time. A wedge of 200° of data were collected using 1.0° per frame strategy. The data were collected on a MarCCD 300 detector at the SER-CAT beamline at the APS. The data were processed and reduced with HKL2000 [16]. The structure (Form C) was initially solved by molecular replacement using one chain of the *Mt*FBPaseII structure (PDB entry 6ayu) yielding two separate chains in the asymmetric unit, but refinement was not pursued at that time.

### Active site and enzyme mechanism modeling

The basis for the modeling of the active site and subsequent hypothesis for the enzyme mechanism was the refined Form A structure (7txg) containing a tetramer in the asymmetric unit. Given the variations within the four chains in the tetramer, chain C was chosen because it had the most complete solvent coordination sphere around the Mn^2+^ native metal cofactor with all the water molecule B-factors in the 30-40 Å2 range, comparable to the surrounding protein atoms (**Fig. SI1**). To position and orient the substrate molecule (F1,6BP) in the active site, the structures of *Ec*FBPaseII (PDB entry 3d1r) and *Mt*FBPaseII (7txb, this work) were superposed. As described (**Fig. 1** and **Fig. SI2**), small differences were observed in the position of the anchoring phosphate (Phosphate-6) and only a minor rotation was needed for the orientation of the oxygens of the cleavable phosphate (Phosphate-1) (see Results section ***Mt***(**Mg^2+^**)**FBPaseII-F1,6BP complex** (**Form C**)).

**Figure 1.**
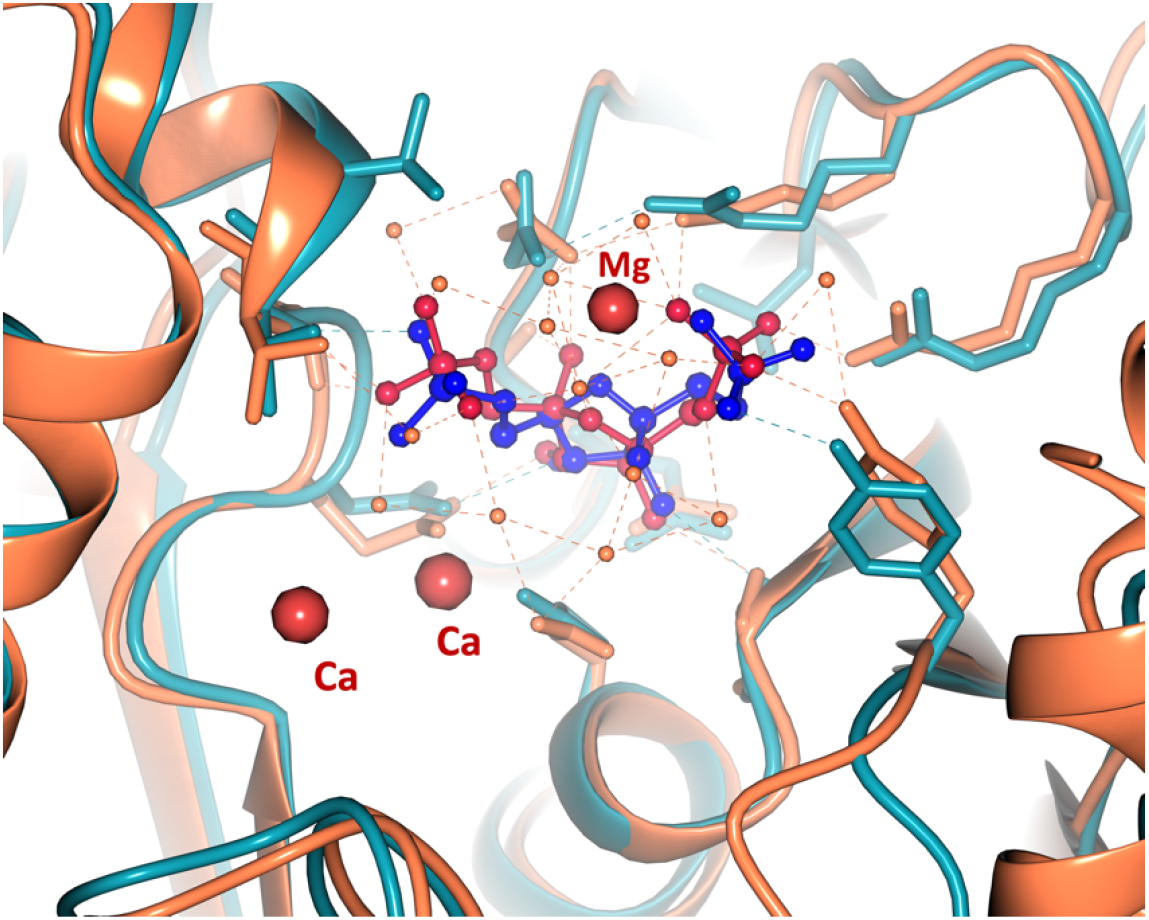
Superposition of F1,6BP (blue) in *Mt*FBPaseII Form C (cyan) and F1,6BP (crimson) in *Ec*FBPaseII (coral red) (PDB 3d1r). Red large spheres mark the position of the metals in the *E. coli* structure.

The position of latter phosphate group displaced several water molecules present in the previously reported structures of *Mt*FBPaseII complexed with the product F1,6P (PDB entries 6ayu, 6ayv). The hydrogen bonds between this phosphate and the N-terminal amide protons of the helical turn G88-T89-T90-I91 were apparent. These minor adjustments revealed the side-chain hydroxyl group of T89 as the most likely candidate for the interaction with the ‘activated water’ (W1), via W2 within the coordination sphere of Mn^2+^. The resulting interatomic distances among the nearest atoms to the Mn^2+^ and relevant amino acids have been illustrated below (**Fig. 5**) within one-tenth of an Ångstrom. The estimated maximum-likelihood coordinate error based on the final refinement parameters at this resolution is 0.20 Å.

## Results and Discussion

### *Ft*FBPaseII and *Mt*FBPaseII activity at various Mn^2+^ and Mg^2+^ concentrations

Initial characterization of the divalent metal cation requirements for the *Mt*FBPaseII enzyme suggested that at low (<3 mM) Mn^2+^ concentrations the enzyme could exhibit a ‘burst of activity’ but high concentrations would significantly inhibit the activity (**Fig. 2A**). The binding of the genuine cofactor Mg^2+^ exhibit the hyperbolic profile of activity vs. concentration upon saturation of the metal binding site(s). In contrast, for *Ft*FBPaseII the data show (**Fig. 2B**) that activity of *Ft*FBPaseII stays high (1.25 ± 0.25 AU) at Mn^2+^ concentrations ranging from 1 mM to 50 mM, whereas same concentrations of Mg^2+^ cause a complete loss of activity. These results complement previous studies of *Ft*FBPaseII activity conducted in a buffer lacking KCl, where a gradual increase in activity of the enzyme was observed over a similar range of concentrations of Mn^2+^ with a maximum at 50 mM [7]. Mg^2+^ was found to be inhibitory at any concentration tried regardless of KCl absence or presence, supporting the notion that the structure of *Ft*FBPaseII complexed with Mg^2+^ that had been reported earlier is that of the inactive enzyme [6].

**Figure 2.**
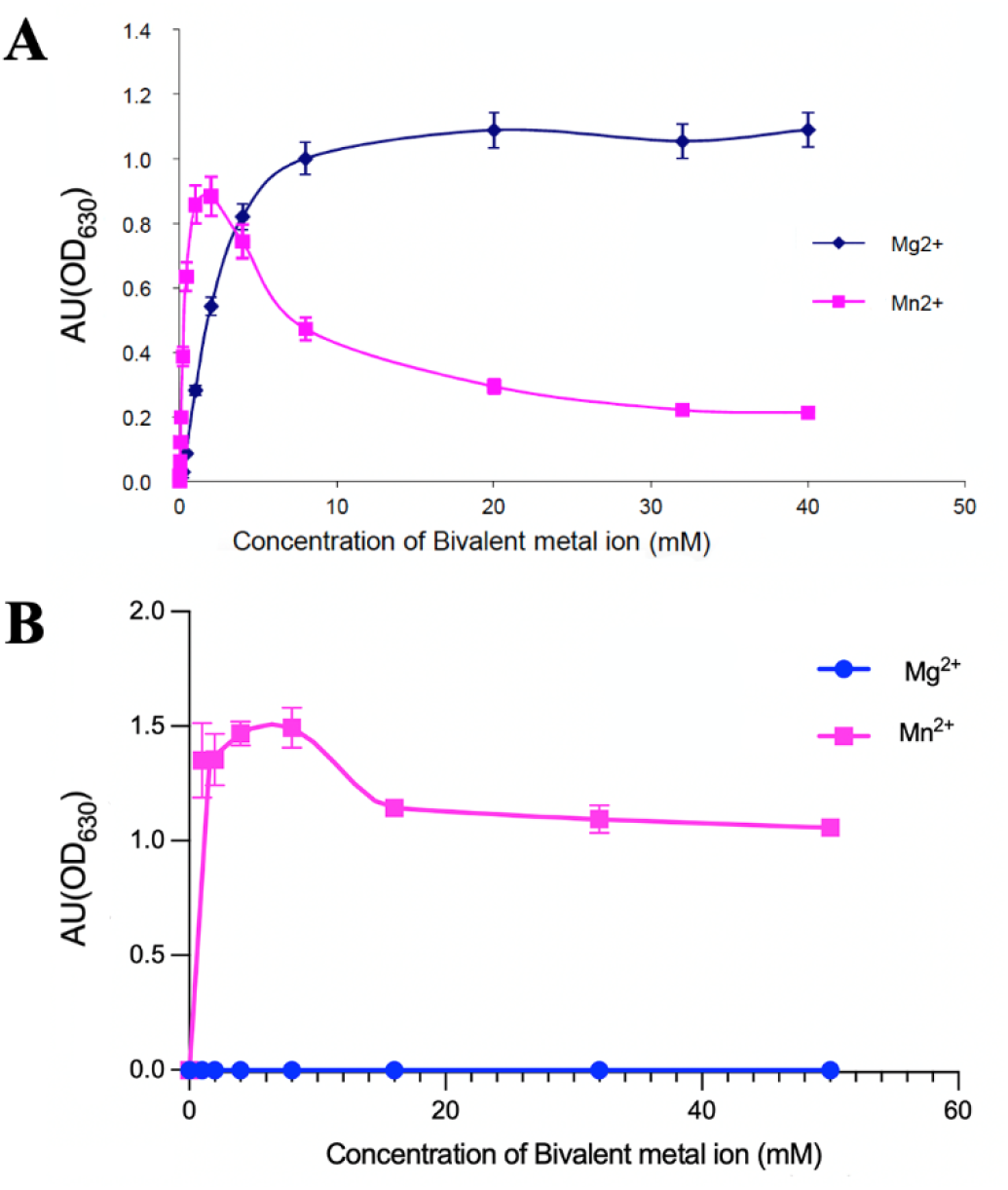
Dependence of *Mt*FBPaseII (**A**) and *Ft*FBPaseII (**B**) activities on the concentration of divalent cations Mn^2+^ and Mg^2+^.

A similar ‘burst of activity’ in the presence of low (~2 mM) concentrations of Mn^2+^ has been observed in the activity of the dual Fructose-1,6/sedoheptulose-1,7-bisphophatase (FBP/SBpase) from the cyanobacteria *Synechocystis* (PCC6803) [8]. Synchronous fluorescence and fluorescence quenching studies on the binding of metal ions were used in combination with experimental binding studies to compare the affinities and fingerprints of various metal cations (Mg^2+^, Mn^2+^, Zn^2+^, Ca^2+^) upon enzyme binding. The results independently confirmed the preference for Mg^2+^ and Mn^2+^ over the other two and showed a similar profile to the one shown in **Fig. 2A**, upon variation of the divalent metal concentration. Estimation of the free energy of binding (ΔG) from the slope of the van’t Hoff plots showed a significant difference between the binding of Mg^2+^ (−22.8 kJ mol^−1^) vs Mn^2+^ (−15.5) at T=295°. More significantly, the binding of Mn^2+^ had a rather large and positive entropy value (205.5 J mol K^−1^) when compared with Mg^2+^ (−467.6 J mol K^−1^) [8]. These quantitative results strongly support the notion that the native divalent cation for PCC6803 is Mg^2+^. In view of the results presented for *Mt*FBPaseII here and the previous structures (6ayy, 6ayu, 6ayv), we hypothesize that the binding of Mn^2+^ for *Mt*FBPaseII at low concentration is transient and does not represent the native cofactor binding in the active site. Conversely, the binding of the non-native Mg^2+^ observed in *Ft*FBPaseII [6] represented a fully inactive conformation of the enzyme. We hypothesize that similar ‘burst of activity’ effects at low concentration of non-genuine metal cofactors in this class of enzymes, could mask and make it difficult to fully characterize the true, genuine, metal dependence profile of class II FBPases and more detail studies are recommended for each member of the class.

### Structure solutions and refinement

#### Form A

Indexing of the initial diffraction snapshots suggested that this crystal form was triclinic (P1) with the following cell parameters a = 64.18, b = 76.23, c = 77.92 Å; α = 68.02°, β = 68.22°, γ = 76.65° diffracting to about 1.9 Å. The size of the unit cell for a P1 lattice suggested that the unit cell of the crystal contained a full tetramer of *Ft*FBPaseII, each chain containing 328 amino acids plus an N-terminal His-tag). Throughout the refinement there were significant differences in the quality and definition of the four different chains near the environment of the metal site waters and active site since no non-crystallographic symmetry (NCS) restraints were used. Statistics for the integration, data reduction, and refinement are presented in **Table SI1**.

#### Form B

Indexing of the initial diffraction snapshots suggested that the crystal form was also P1 with cell parameters a = 65.82, b = 76.37, c = 141.11 Å; α = 76.98°, β = 87.10°, γ = 75.84° and diffracting only to about 2.4 Å. These crystals were more radiation sensitive than those of form A and thus a compromise between data quality and exposure time was adopted. The size and symmetry of the unit cell, when compared to form A, suggested that the crystal cell contained two tetramers of *Ft*FBPaseII (328 amino acids plus an N-terminal His-tag). The conformation of most external loops varied quite significantly among different chains, possibly explaining the low symmetry of both crystal forms. Statistics for the integration, data reduction and refinement are presented in **Table SI1**.

#### Form C

An initial analysis of these crystals showed that they were the same hexagonal space group P6_1_22 as the previously analyzed *Mt*FBPaseII crystals [19] (PDB entries 6ayy, 6ayu, 6ayv) but with an approximately double c axis (a = b = 117.1, c = 325.92 Å, vs. c = 140 Å) diffracting only to about 3.7Å. The interest on these crystals was renewed when a preliminary Molecular Replacement (MR) solution of the structure suggested the presence of the substrate F1,6BP in a very loose packing arrangement of *Mt*FBPaseII tetramers. The initial MR solution was later confirmed, and the structure was fully refined. Given the limited resolution of the data, the refinement proceeded initially with NCS restrains as the main aim of the analysis was to provide a robust electron density to fit the substrate in the active site pocket of the two chains in the asymmetric unit. Later the NCS restrains were removed and refinement proceeded with conventional protocols. Statistics for the integration, data reduction, and refinement are presented in **Table SI1**.

#### Overall view of the structures *Ft*(Mn^2+^)FBPaseII-Glycerol/F6P complex (Form A)

The high resolution (1.9 Å) for this structure revealed one approximate 222 (D_2_) tetramer in the triclinic cell. The overall quaternary arrangement of the four chains did not differ significantly from the tetramer described earlier for the Mg^2+^ liganded structure (PDB entry 7js3). However, significant local differences were observed in chains B and D, particularly in the residues related to the orientation and disposition of the protruding helix (H12, Chain C Phe235-Leu252; Chain D Ala231-Leu261) (**Fig. 3**). Both involved in crystal contacts on this triclinic form. The tertiary structures did not differ significantly from the previously reported structure however chains A and B showed weak density for the residues in the protruding helix Ala234-Ala248 and chain D in the external residues Gly56-Met65 of the Ψ loop [8]. All the mentioned regions also showed significant local deviations from the NCS using subunit A as a reference.

**Figure 3.**
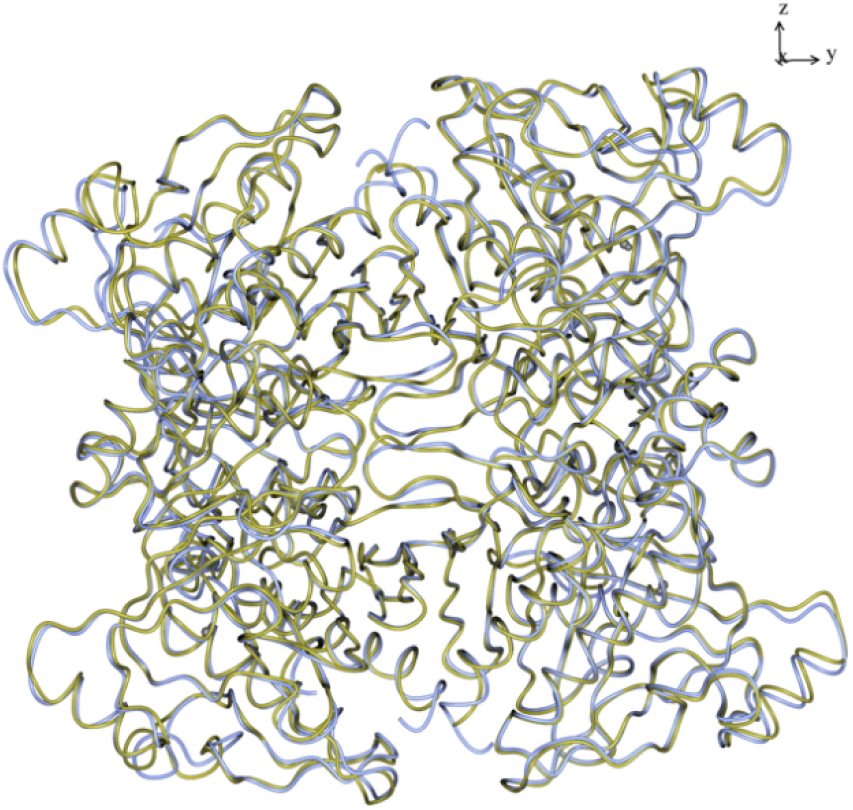
Superposition of the D_2_ tetramer of the *Ft*(Mg^2+^)FBPaseII (PDB 7js3) (gold) and the corresponding oligomer of *Ft*(Mn^2+^)FBPaseII-F6P, Form A (ice blue). The relative orientation of the four chains in the tetramer is conserved and only relatively minor local displacements are observed in the outer most loops of the tetramer. The inhibition of the *Ft*FBPaseII by the Mg^2+^ cation does not affect the overall structure of the tetramer.

Several discrete glycerol molecules were refined to reasonable B factors (<80 Å^2^) at distinct locations in the four chains A(1), B(5), C(2), and D(4). The phosphate groups were refined at the putative locations of the product F6P (residues Arg165-Pro-Arg167) in chains B, C, and D. Attempts to refine full F6P molecules at these locations failed because of the presence of well refined glycerol molecules that apparently competed with the full F6P occupancy (**Fig. 4**).

**Figure 4.**
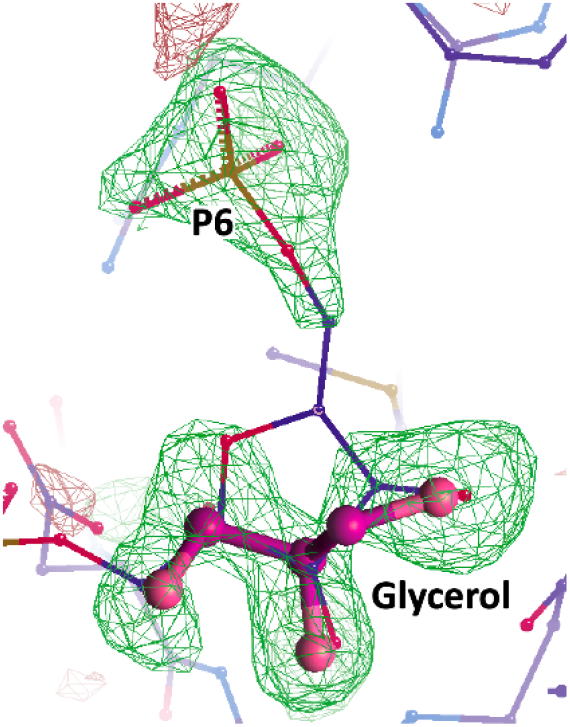
Omit electron density map of the refined structure of *Ft*FBPaseII (Chain C) centered on the product binding region showing the partial occupancy of F6P and the corresponding fitting of a bound glycerol molecule.

A chemically consistent environment for the Mn^2+^ cofactor in this crystal structure was refined in the proximity of the F6P for chain C (**Fig. 5** and **Fig. SI1**). The structure revealed an octahedrally coordinated Mn^2+^ ion, surrounded by the side chain carboxylate group of Asp84, the main chain carbonyl oxygen of Leu86 and four water molecules. In addition, the side chain of Glu57 projects its carboxylate group towards the lower apical liganded water (distance 3.0 Å) and could play a role in the coordination of the metal since this portion of the chain is disordered in crystals of *Ft*FBPaseII grown in the absence of metal cations (Mg^2+^ or Mn^2+^, unpublished observations).

**Figure 5.**
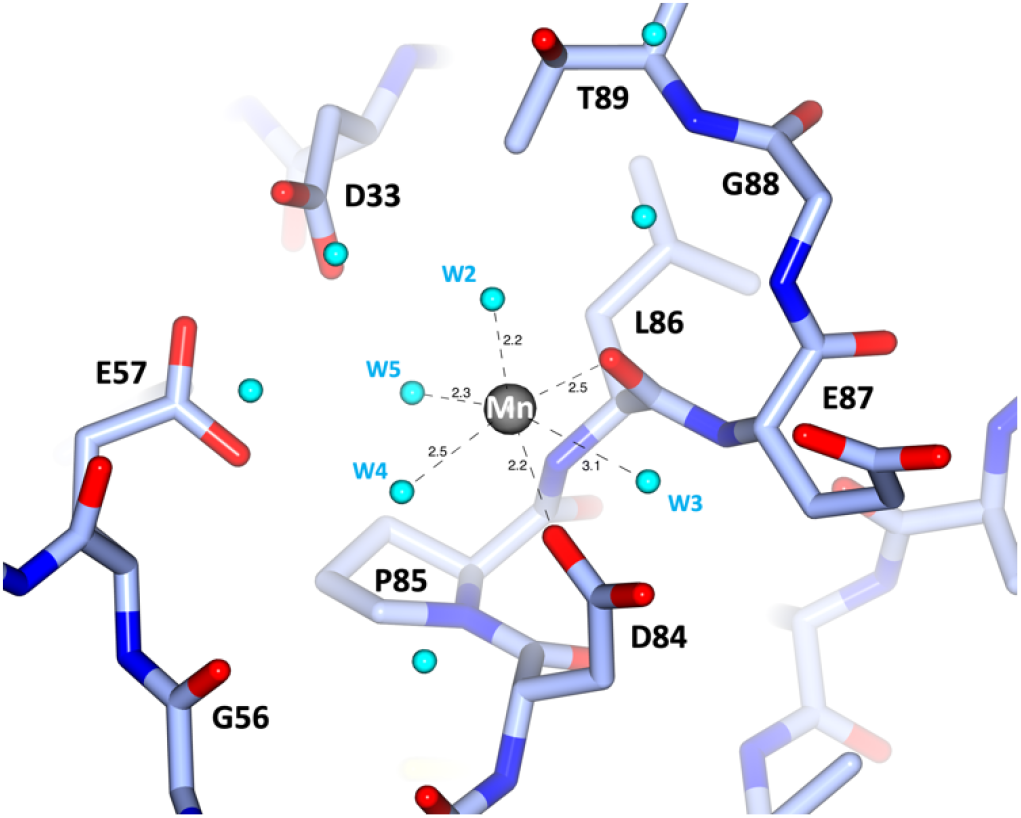
Distances and residues involved in the coordination of Mn^2+^ in the best-defined (chain C) active site of Form A. See the corresponding electron density map in **Fig. SI1**.

#### *Ft*(Mn^2+^)FBPaseII-F6P complex (Form B)

Form B represents the most complex crystallographic structure of *Ft*FBPaseII observed to date. The triclinic cell contained two approximate 222 (D_2_) tetramers separated by about 70 Å with the orientation of their respective local dyads being slightly different. The overall structures of the two tetramers did not differ substantially as compared to the tetramer observed in the Mg^2+^ containing structure (PDB 7js3) although — as discussed before for the Form A crystals — there were local differences in the external loops involved in the crystal contacts. The most striking observation of this crystal form was the presence of the product of the reaction F6P in only one subunit of each of the tetramers. This is illustrated in **Fig. 6A**. Surprisingly, the position and orientation of the product is somewhat displaced from the position observed in *Mt*FBPaseII complexes (PDB 6ayu, 6ayv) (**Fig. 6B**). A possible interpretation of these observations is discussed later.

**Figure 6.**
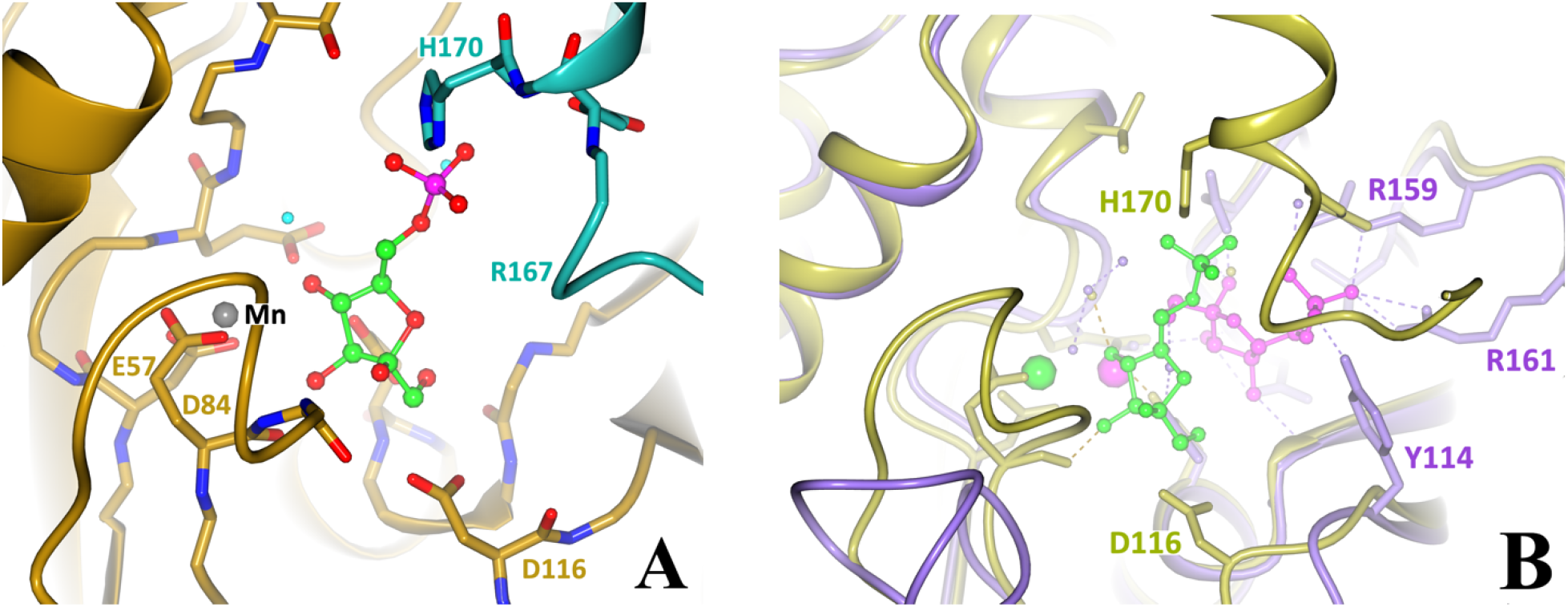
**A.** Molecule of F6P found in the crystal contact area of one chain each of the two tetramers in the unit cell of the P1 crystals of *Ft(*Mn^2+^)FBPaseII-F6P (Form B). The two subunits making the crystal contact are depicted in different colors. Surrounding residues are labeled and the Mn^2+^ is shown as a grey sphere. **B.** Superposition of F6P (magenta) complexed with *Mt*FBPaseII (purple) in the ‘canonical’ (previously observed) binding of the product F6P (PDB 6ayv), compared with binding position of F6P (green) observed in the crystal contacts of the *Ft*FBPaseII (Form B) structure (gold). Lower left demonstrates the local conformational change taking place in the P1 crystal lattice (gold) vs. the hexagonal crystals of *Mt*FBPaseII (purple). Green and magenta spheres mark the position of Mn^2+^ and Mg^2+^ cations observed.

#### *Mt*(Mg^2+^)FBPaseII-F1,6BP complex (Form C)

The MR solution of this crystal form placed two separate chains (A, B) in the crystal asymmetric unit. The final refined structure confirmed the presence of the F1,6BP substrate in both chains and was used as a reference in conjunction with the *E. coli* structure (PDB 3d1r) to model the position and orientation of the substrate in the active site. The position of the substrate phosphate groups between the two structures differed only by approximately 0.7 Å (**Fig. 1** and **Fig. SI2**) and was considered in reasonable agreement to support the proposed enzymatic mechanism for *Mt*FBPaseII, *Ec*FBPaseII, *Ft*FBPaseII and homologous class II FBPases, given the lower resolution of the first structure (see Supplementary Information Table 1). The two separate chains appear to have a Mg^2+^ cation bound partially by the phosphate-6 of the substrate, but the resolution of the structure does not permit any further inferences. In the final stages of refinement, it was possible to reassemble the asymmetric unit of the crystal to form a dimer of the two separate chains, homologous to the one suggested in the initial *E. coli* structure but different from the one found in the asymmetric unit of the PDB entry 6ayu. The two different combined dimers form the consensus tetrameric structure of the class II FBPases that is formed by crystallographic symmetry as discussed previously [19]. All four sites were occupied by the substrate F1,6BP. We hypothesized that the observation of the uncleaved substrate in this crystal form was due to the limited availability of the native divalent cation (Mg^2+^) since the concentration was only 1 mM in the crystallization media and the crystal structures do not provide enough evidence for full occupancy of the sites (**Fig. 2A**).

#### Glycerol molecules

In addition to the water molecules found in the two high resolution structures of *Ft*(Mn^2+^)FBPaseII presented here (Form A: 7txg, Form B: 7txa) (**Table SI1**), it is worth reviewing several glycerol molecules found in the highest resolution structure of the two (7txg, **Table SI2**). A total of 11 glycerol molecules with robust refinement parameters and acceptable temperature factors <90 Å were found. Not surprisingly, the majority of them were found in chains C and D that are the chains with lower overall temperature factors and best-defined electron density. The glycerol molecules with lowest B-factors (B<40 Å^2^) are: C701, C601, D501, D403, and D502. The first one is competing with the lower part of the F6P molecule in the active site of chain C (**Fig. SI3**) and will be discussed later.

Of the remaining glycerol molecules, the most relevant is D403. The quality of the density and surface rendering of the pocket is illustrated in (**Fig. SI3**). This ‘site’ is occupied by a homologous glycerol molecule in the structure of the previously reported *Ft*(Mg^2+^)FBPaseII (PDB entry 7js3) (**Fig. 7**).

**Figure 7.**
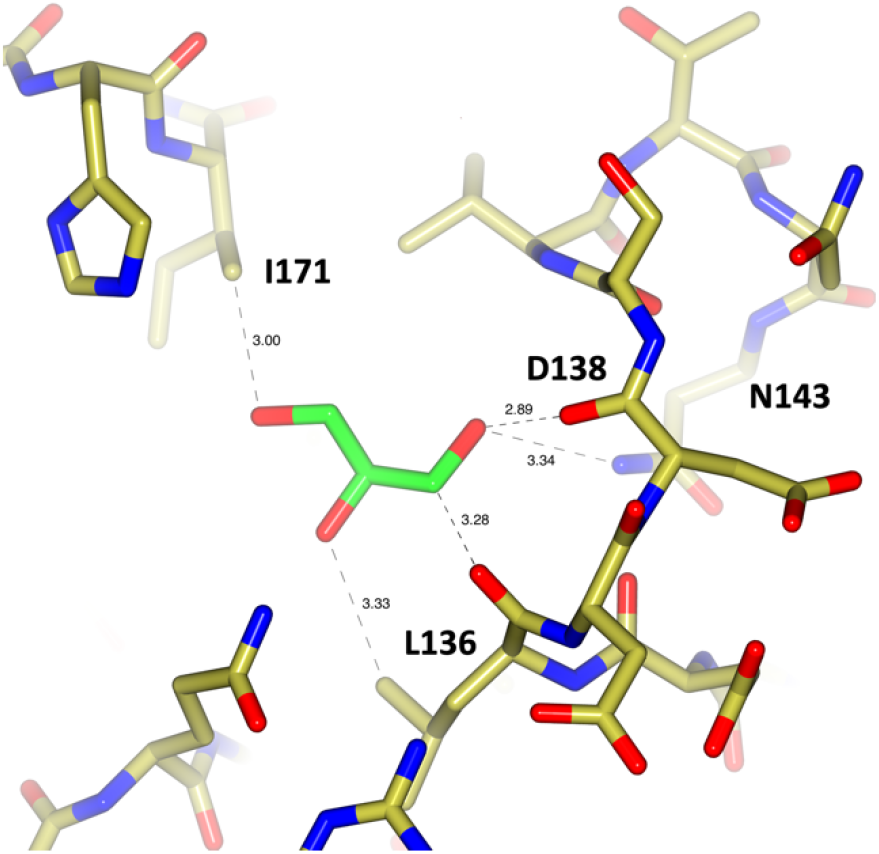
*Ft*(Mg^2+^)FBPaseII (PDB 7js3). Amino-residues surrounding Glycerol D403 in the structure of *Ft*(Mg^2+^)FBPaseII within a cutoff distance of 3.4 Å.

Analogous glycerol molecules have been found also in the number of *Mt*FBPaseII structures [19] (PDB entries: 6ayy, 6ayu, 6ayv). In all the structures, the pocket is formed by a five residue ‘arch’ over the N-terminal Ψ-loop that in *Ft*FBPaseII includes residues L136-D137-D138-S139-V140 (*Mt*FBPaseII: I130-T131-A132-P133-I134). Most of the interactions of the glycerol hydroxyl groups are with main chain atoms of the polypeptide chain. (**Fig. 7** and **Fig. SI3**). Arg232 (*Ft*) is partially covering the entrance to the pocket and provides the positively charged surface highlighted in **Fig. SI3**. The observed glycerol pocket is covered by the last α-helix layer of the five-layer α-β-α-β-α sandwich (residues 122-180 including the Arg169-Pro170-Arg171) that is characteristic of the structure of this class of enzymes. There is a short external solvent channel from the glycerol site to the guanidinium group of Arg171 in the active site, which participates in anchoring the Phosphate-6 of the substrate F1,6BP.

### Catalytic mechanism

A novel catalytic mechanism behind the hydrolysis of F1,6BP by class II FBPases is being proposed and discussed here. The activity of all class II FBPases has an absolute requirement for one or more divalent cations, depending upon the organism. Our crystallographic data identified a single Mn^2+^ ion in the active site 7.2 Å from Phosphate-1 and held in an octahedral coordination with four water molecules, the carbonyl main chain of L86 and the carboxylate side chain of D84. Modeling the position of the missing phosphate group in the structure of *Ft*FBPaseII liganded with F6P enabled the estimation of the distances to the cleavable Phosphate-1 group and the identification of atoms that could coordinate it during catalysis (**Fig. 8A**). This led to the observation that the N-terminus of a two turn α-helix (H3; G88-T89-T90-I91-T92-S93-K94) is facing Phosphate-1, about 3.1 Å away.

**Figure 8.**
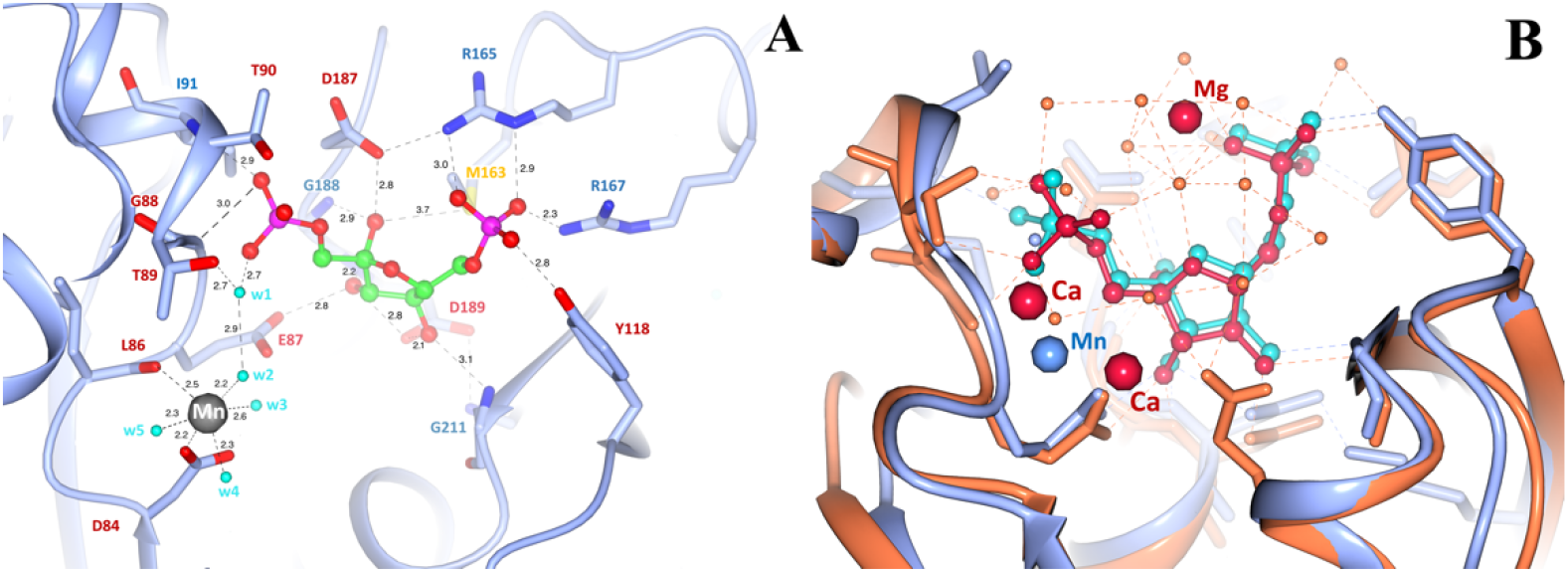
**A.** Active state and metal coordination in *Ft*(Mn2+)FBPaseII (7txg). **B.** Superposition of modeled F1,6BP (aqua) in *Ft*FBPaseII Form A (ice blue) with F1,6BP (crimson) found in *Ec*FBPaseII (PDB 3d1r) (coral red). Annotated distances are from the refined structure as described in “Materials and Methods”.

Based on these observations, we propose a 1-metal/phosphate-binding helix assisted catalytic mechanism as follows: The Mn^2+^ ion positions and activates two water molecules (W1 – indirect, W2 - direct coordination; **Figs 8-10**) via stabilizing the formation of hydroxide anions for nucleophilic addition to Phosphate-1. In addition, our observations and proposed mechanism are fully consistent with the previously demonstrated critical role of the hydroxyl sidechain of Thr89 in the catalytic mechanism, where the T89S variant of *Ft*FBPaseII is active (albeit slower), but the T89A is completely inactive [8]. The T89/S89 amino acid residue is critical for catalysis but does not provide any significant binding affinity for the substrate. Mutation of the homologous residues of the *Mt*FBPaseII enzyme exhibit similar enzymatic properties (T84S is partially active but T84A is completely inactive) [19]. A possible structural explanation of this effect can be surmised based on the additional steric hindrance of the sidechain of the Thr residue provided by the neighboring residue Leu86 (Ile87 in *E. coli*), facilitating the correct positioning and orientation of the critical hydroxyl group (**Fig. 9**).

**Figure 9.**
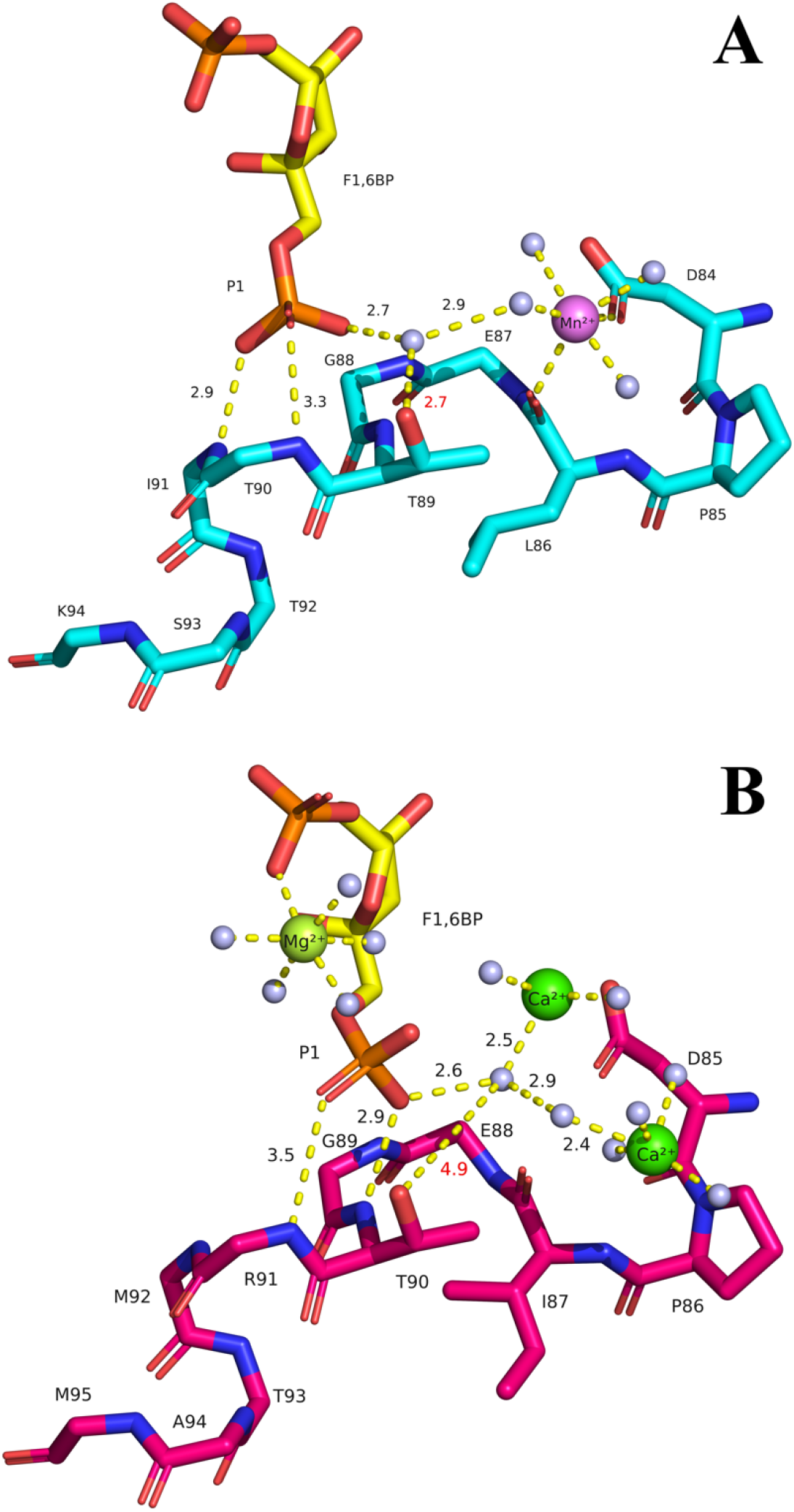
Proposed stabilization of the leaving phosphate group (P1) during the catalytic process for FBPases class II. Ligands coordination by active *Ft*FBPaseII (**A**) and inactive *Ec*FBPaseII (PDB 3d1r) (**B**). The interatomic distances and the side-chains important amino acid residues have been highlighted. Waters are shown as blue spheres. Note the hydrogen bond interactions of the leaving phosphate group (P1) with the amide protons of the N-terminal helix in both structures. The presence of L86 (I87 in *E. coli,* **B**) probably orients and positions optimally the side chain OH group of T89 for the catalytic action. The T89S mutation would result in a less restrained OH yielding an enzyme with a lower catalytic activity; the T89A mutation resulted in a totally inactive enzyme [6].

**Figure 10.**
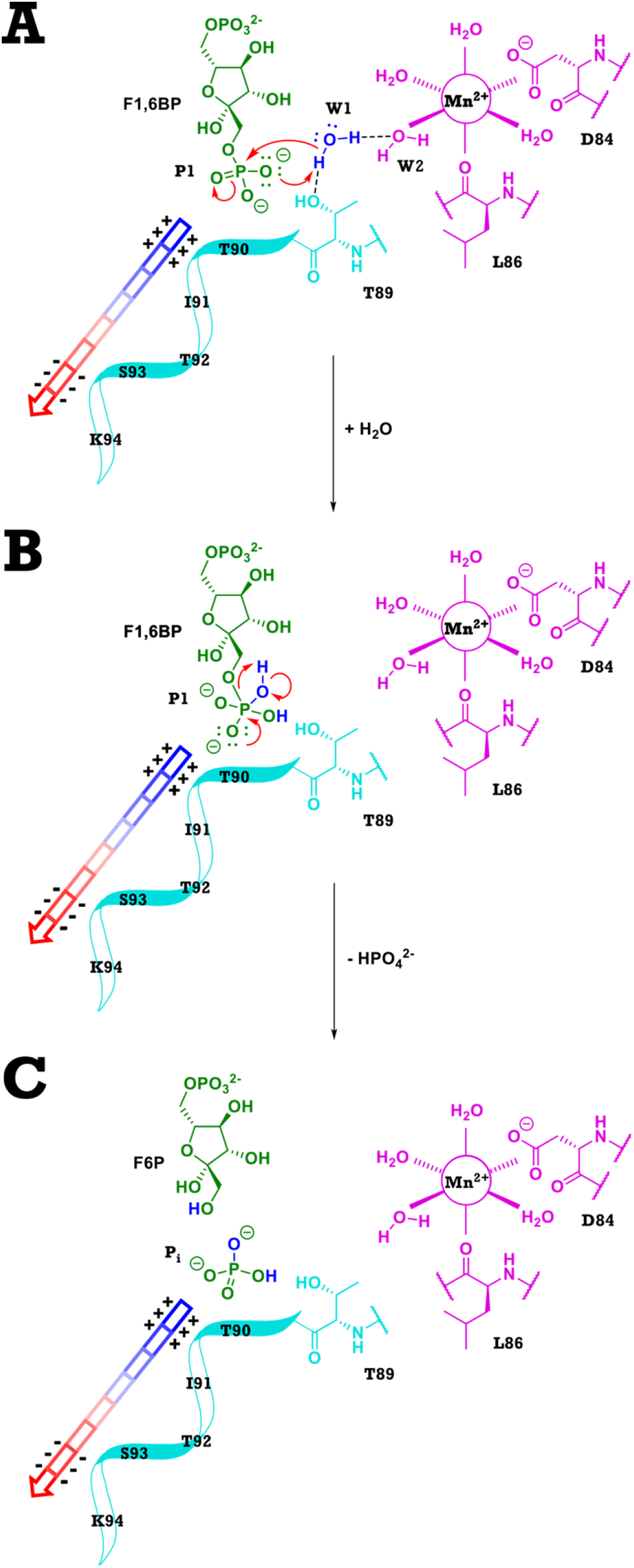
Details of the proposed catalytic mechanism. **A.** Addition/attack of W1 to phosphate-1 of F1,6BP. **B.** Elimination of phosphate intermediate by cleaving P-O bond. **C.** Product of hydrolysis yields F6P and an inorganic phosphate. The stabilization of the transition state intermediate (**B**) and of the leaving PO^2-^_4_ group is accomplished by the positive charge (blue) of the helix N-terminal dipole indicated by the arrow.

The phosphate binding helix assists in the binding of F1,6BP in the active site as well as stabilizes the negatively charged tetrahedral intermediate transition states of Phosphate-1 after the nucleophilic addition of water. The helix further decreases the rate of the reverse reaction by coordinating the inorganic phosphate upon elimination, (**Fig. 9–10**). This helix dipole moment arises because the approximate 3.5 Debyes dipole moment from the formal charges of the amide oxygen and nitrogen in each individual peptide bond neatly align along the helix axis. The strength of the field is proportional to the length of the helix but tapers after two turns which is the size of the helix found in the active site [20]. This is not unusual as the enzymes lactate dehydrogenase, glyceraldehyde phosphate dehydrogenase and triosephosphate isomerase all contain phosphate binding helices that bind to the negatively charged phosphate groups of NAD and/or glyceraldehyde-3-phosphate at the N-terminus. The dipole moment has the advantage of having the effect of an isolated charge without being excessively solvated or in proximity of a salt bridge, as is typical for charged residues [20]. A schematic representation of the catalytic mechanism is presented in **Fig. 10**. The helical structure highlighted above is conserved in all FBPasesII for which the three-dimensional structure is known and the amino acid sequences, if not fully conserved (**Fig. 11**), do not have any insertions or deletions in that region that could indicate significant disruptions of the helical structure, as shown for *E. coli* and *F. tularensis* (**Fig. 9**).

**Fig. 11.**
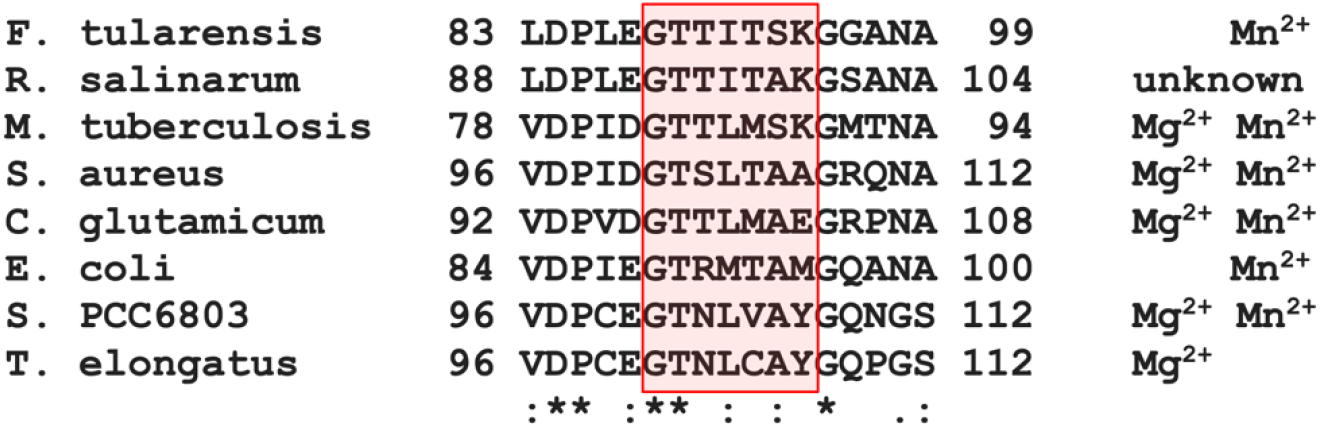
Sequence alignment of several class II FBPases. The selected region includes the highly conserved critical Thr89 (*Ft*FBPaseII) residue and the following two-turn helix whose helical dipole is suggested to be implicated in the catalytic mechanism. Divalent cations required for activity are listed.

The detailed number and position of the divalent cations in the active Mg^2+^-dependent FBPasesII such as *Mt*FBPaseII, as well as their coordination spheres, is still uncertain in view of the limited value of the available PDB structures (6ayy, 6ayu, 6ayv and homologous entries) as indicated in the introduction. However, the structural insights derived from this work strongly suggests that activated water(s) molecules via the divalent cation(s) (Mn^2+^or Mg^2+^) combined with the hydroxyl group of Thr89 (or homologous e. g. Ser89) residue would be required for catalysis in all class II FBPases. In addition, it is reasonable to assume that the conserved N-terminal helical turn dipole would also play a role in the stabilization of the transition phosphate intermediate in the Mg^2+^-dependent subgroup of FBPasesII, based on the conservation of the helical feature (**Fig. 11**). Further structural and biochemical studies on other members of the class II FBPase enzymes are planned to confirm these generalizations.

## Conclusions

We have been able to infer a consistent binding mode of the F1,6BP substrate in the active site pocket of class II FBPases in the presence of the native metal cofactors and surrounding water molecules. This was accomplished by analysis of the available structural information from previous publications and PDB deposited structures of class II FBPases from *E. coli*, *M. tuberculosis* and *F. tularen*sis, combined with the three additional crystal structures presented here. These include active *Ft*FBPaseII complexed with a native metal cofactor Mn^2+^ and F6P product, together with the structure of substrate bound *Mt*FBPaseII. This allowed us to propose a catalytic mechanism for the entire class II of FBPases, consistent with conserved structural features in the proximity of the active site for the enzymes of the three species used here for analysis. Namely, the presence of a critical hydroxyl-containing residue (Thr), and the existence of an N-terminal helical turn that provides a dipole-induced positive charge to stabilize the transition state of the cleavable Phosphate-1 (leaving group). Although the suggested catalytic mechanism has not been proven in this work, the proposed mechanism is more consistent chemically and structurally, than the earlier proposals based on the initial *E. coli* structure (PDB 3d1r). We hypothesize that this mechanism will also be applicable to other members of the class based on the high conservation of the amino acid sequences around those structural features. The specific native divalent cation may vary, most commonly Mg^2+^ or Mn^2+^, but the critical water coordination sphere around the native divalent cation, and the role of the conserved helical dipole would remain the key structural features of the enzymatic mechanism.

## Abbreviations used

FBPase: fructose-1,6-bisphophatase
F1,6BP: fructose 1,6 bisphosphate
F6P: fructose-6-phosphate
*Ft*FBPase: *F. tularensis* FBPase
*Mt*FBPase: *Mycobacterium tuberculosis* FBPase
*Ec*FBPase: *E. coli* FBPase
*Ft*FBPaseII: class II *Ft*FBPase and similarly for the others.
*Ft*(Mg^2+^)FBPase: *F. tularensis* FBPase containing a Mg^2+^ divalent cation or crystallized in the presence of such a cation
*Ft*(Mn^2+^)FBPase: *F. tularensis* FBPase containing a Mn^2+^ divalent cation or crystallized in the presence of such a cation
NAD: nicotinamide dinucleotide phosphate
PDB: Protein Data Bank.

## Associated content

**Supporting information**

**Table SI1.** Data reduction and refinement statistics for the three crystal structures.

**Table SI2.** Ligands in *Ft*(Mn^2+^)FBPase-F6P (7txg).

**Figure SI1.** Mn^2+^ coordination in the active site of Chain C in the structure of *Ft(*Mn^2+^)FBPaseII-Glycerol/F6P, Form A (Chain C). 2F_o_-F_c_ electron density map contoured at 1.5σ

**Figure SI2.** Omit (F_o_-F_c_) electron density map of the refined structure of *Mt*FBPaseII (Form C).

**Figure SI3.** *Ft*(Mn^2+^)FBPase (7txg).

## Author information

**Corresponding authors**

Farahnaz Movahedzadeh - Institute for Tuberculosis Research, University of Illinois at Chicago, Chicago, Illinois, USA and Department of Pharmaceutical Sciences, University of Illinois at Chicago, Chicago, Illinois, USA.

Celerino Abad-Zapatero - Institute for Tuberculosis Research, University of Illinois at Chicago, Chicago, Illinois, USA and Center for Biomolecular Sciences, University of Illinois at Chicago,

## Authors

Anna I. Selezneva - Institute for Tuberculosis Research, University of Illinois at Chicago, Chicago, Illinois, USA

Luke N.M. Harding - Department of Pharmaceutical Sciences, University of Illinois at Chicago, Chicago, Illinois, USA

Hiten Gutka - Institute for Tuberculosis Research, University of Illinois at Chicago, Chicago, Illinois, USA. Current address: Bristol-Myers Squibb, Summit, New Jersey, USA.

## Authors contributions

**Anna I. Selezneva:** Conceptualization, Investigation, Data curation, Methodology, Visualization, Writing - Original Draft, Review & Editing, Validation.

**Luke N. M. Harding:** Investigation, Visualization, Writing - Original Draft, Writing-Review & Editing. Hiten Gupta: Resources.

**Farahnaz Movahedzadeh:** Supervision, Project administration, Review & Editing, Funding acquisition.

**Celerino Abad-Zapatero:** Conceptualization, Methodology, Investigation, Visualization, Writing - Original Draft, Review & Editing, Supervision, Data Curation, Funding acquisition.

## Notes

### Data availability – PDB accession codes

*Ft(*Mn^2+^)FBPaseII-Glycerol/F6P (Form A) – 7txg

*Ft(*Mn^2+^)FBPaseII-F6P (Form B) – 7txa

*Mt*(Mg^2+^)FBPaseII-F1,6BP complex (Form C) – 7txb

## Acknowledgements

The Advanced Photon Source was supported by the US Department of Energy (DE-AC02-06CH11357). We acknowledge the help of the LS-CAT staff at the Advanced Photon Source, Argonne National Laboratory with data collection on the LS-CAT 21-ID beamlines. The authors would also like to thank Professor Scott Franzblau for his support to this project under the auspices of the Institute of Tuberculosis Research.

## References

1. Kadzhaev K, Zingmark C, Golovliov I, Bolanowski M, Shen H, Conlan W, et al. Identification of genes contributing to the virulence of Francisella tularensis SCHU S4 in a mouse intradermal infection model. PLoS One. 2009;4(5):e5463. Epub 2009/05/09. doi: 10.1371/journal.pone.0005463. PubMed PMID: 19424499; PubMed Central PMCID: PMCPMC2675058.

2. https://emergency.cdc.gov/agent/agentlist-category.asp#a.

3. Santic M, Molmeret M, Klose KE, Abu Kwaik Y. Francisella tularensis travels a novel, twisted road within macrophages. Trends Microbiol. 2006;14(1):37–44. Epub 2005/12/17. doi: 10.1016/j.tim.2005.11.008. PubMed PMID: 16356719.

4. Brown G, Singer A, Lunin VV, Proudfoot M, Skarina T, Flick R, et al. Structural and biochemical characterization of the type II fructose-1,6-bisphosphatase GlpX from Escherichia coli. J Biol Chem. 2009;284(6):3784–92. Epub 2008/12/17. doi: 10.1074/jbc.M808186200. PubMed PMID: 19073594; PubMed Central PMCID: PMCPMC2635049.

5. Donahue JL, Bownas JL, Niehaus WG, Larson TJ. Purification and characterization of glpX-encoded fructose 1, 6-bisphosphatase, a new enzyme of the glycerol 3-phosphate regulon of Escherichia coli. J Bacteriol. 2000;182(19):5624–7. Epub 2000/09/15. doi: 10.1128/JB.182.19.5624-5627.2000. PubMed PMID: 10986273; PubMed Central PMCID: PMCPMC111013.

6. Selezneva AI, Gutka HJ, Wolf NM, Qurratulain F, Movahedzadeh F, Abad-Zapatero C. Structural and biochemical characterization of the class II fructose-1,6-bisphosphatase from Francisella tularensis. Acta Crystallogr F Struct Biol Commun. 2020;76(Pt 11):524–35. Epub 2020/11/03. doi: 10.1107/S2053230X20013370. PubMed PMID: 33135671; PubMed Central PMCID: PMCPMC7605111.

7. Gutka HJ, Wolf NM, Bondoc JMG, Movahedzadeh F. Enzymatic Characterization of Fructose 1,6-Bisphosphatase II from Francisella tularensis, an Essential Enzyme for Pathogenesis. Appl Biochem Biotechnol. 2017;183(4):1439–54. Epub 2017/05/27. doi: 10.1007/s12010-017-2512-6. PubMed PMID: 28547120; PubMed Central PMCID: PMCPMC5698383.

8. Sun Y, Liao X, Li D, Feng L, Li J, Wang X, et al. Study on the interaction between cyanobacteria FBP/SBPase and metal ions. Spectrochim Acta A Mol Biomol Spectrosc. 2012;89:337–44. Epub 2012/01/17. doi: 10.1016/j.saa.2011.12.014. PubMed PMID: 22244776.

9. Marcus F, Rittenhouse J, Chatterjee T, Hosey MM. Fructose-1,6-bisphosphatase from rat liver. Methods Enzymol. 1982;90 Pt E:352–7. Epub 1982/01/01. doi: 10.1016/s0076-6879(82)90155-0. PubMed PMID: 6296610.

10. Rashid N, Imanaka H, Kanai T, Fukui T, Atomi H, Imanaka T. A novel candidate for the true fructose-1,6-bisphosphatase in archaea. J Biol Chem. 2002;277(34):30649–55. Epub 2002/06/18. doi: 10.1074/jbc.M202868200. PubMed PMID: 12065581.

11. Baykov AA, Evtushenko OA, Avaeva SM. A malachite green procedure for orthophosphate determination and its use in alkaline phosphatase-based enzyme immunoassay. Anal Biochem. 1988;171(2):266–70. Epub 1988/06/01. doi: 10.1016/0003-2697(88)90484-8. PubMed PMID: 3044186.

12. Kimber MS, Vallee F, Houston S, Necakov A, Skarina T, Evdokimova E, et al. Data mining crystallization databases: knowledge-based approaches to optimize protein crystal screens. Proteins. 2003;51(4):562–8. Epub 2003/06/05. doi: 10.1002/prot.10340. PubMed PMID: 12784215.

13. Vagin A, Teplyakov, A.. MOLREP: an Automated Program for Molecular Replacement. J Appl Cryst 1997;30:1022–5. doi: https://doi.org/10.1107/S0021889897006766.

14. Winn MD, Ballard CC, Cowtan KD, Dodson EJ, Emsley P, Evans PR, et al. Overview of the CCP4 suite and current developments. Acta Crystallogr D Biol Crystallogr. 2011;67(Pt 4):235–42. Epub 2011/04/05. doi: 10.1107/S0907444910045749. PubMed PMID: 21460441; PubMed Central PMCID: PMCPMC3069738.

15. McNicholas S, Potterton E, Wilson KS, Noble ME. Presenting your structures: the CCP4mg molecular-graphics software. Acta Crystallogr D Biol Crystallogr. 2011;67(Pt 4):386–94. Epub 2011/04/05. doi: 10.1107/S0907444911007281. PubMed PMID: 21460457; PubMed Central PMCID: PMCPMC3069754.

16. Otwinowski Z, Minor W. Processing of X-ray diffraction data collected in oscillation mode. Methods Enzymol. 1997;276:307–26. Epub 1997/01/01. PubMed PMID: 27754618.

17. Liebschner D, Afonine PV, Baker ML, Bunkoczi G, Chen VB, Croll TI, et al. Macromolecular structure determination using X-rays, neutrons and electrons: recent developments in Phenix. Acta Crystallogr D Struct Biol. 2019;75(Pt 10):861–77. Epub 2019/10/08. doi: 10.1107/S2059798319011471. PubMed PMID: 31588918; PubMed Central PMCID: PMCPMC6778852.

18. Murshudov GN, Skubak P, Lebedev AA, Pannu NS, Steiner RA, Nicholls RA, et al. REFMAC5 for the refinement of macromolecular crystal structures. Acta Crystallogr D Biol Crystallogr. 2011;67(Pt 4):355–67. Epub 2011/04/05. doi: 10.1107/S0907444911001314. PubMed PMID: 21460454; PubMed Central PMCID: PMCPMC3069751.

19. Wolf NM, Gutka HJ, Movahedzadeh F, Abad-Zapatero C. Structures of the Mycobacterium tuberculosis GlpX protein (class II fructose-1,6-bisphosphatase): implications for the active oligomeric state, catalytic mechanism and citrate inhibition. Acta Crystallogr D Struct Biol. 2018;74(Pt 4):321–31. Epub 2018/04/14. doi: 10.1107/S2059798318002838. PubMed PMID: 29652259; PubMed Central PMCID: PMCPMC5892879.

20. Hol WG, van Duijnen PT, Berendsen HJ. The alpha-helix dipole and the properties of proteins. Nature. 1978;273(5662):443–6. Epub 1978/06/08. doi: 10.1038/273443a0. PubMed PMID: 661956.

